# Gateway Identity and Spatial Remapping in a Combined Grid and Place Cell Attractor

**DOI:** 10.1101/2022.05.11.491479

**Authors:** Tristan Baumann, Hanspeter A. Mallot

## Abstract

The spatial specificities of hippocampal place cells, i.e., their firing fields, are subject to change if the rat enters a new compartment in the experimental maze. This effect is known as remapping. It cannot be explained from path integration (grid cell activity) and local sensory cues alone but requires additional knowledge of the different compartments in the form of context recognition at the gateways between them. Here we present a model for the hippocampal-entorhinal interplay in which the activity of place and grid cells follows a joint attractor dynamic. Place cells depend on the current grid cell activity but can also reset the grid cell activity in the remapping process. Remapping is triggered by the passage through a gateway. When this happens, a previously stored pattern of place cell activity associated with the gateway is reactivated from a “gateway database”. The joint attractor will then reinstate the grid cell pattern that was active when the gateway had first been learned and path integration can proceed from there. The model is tested with various mazes used in the experimental literature and reproduces the published results, and we make predictions for remapping in a new maze type. We propose the involvement of memory in the form of “gate cells” that drive the place cells and with them the joint hippocampal-entorhinal loop into the corresponding attractor whenever a compartment is entered.

## Introduction

The neural substrate of spatial representation, the *cognitive map*, is generally thought to be found in place cells located in the hippocampus (HC). First described by O’Keefe & Dostrovsky (1971), these cells fire maximally whenever the animal is within a small localized region in space, the cell’s place field (O’Keefe & Nadel, 1978; Moser et al., 2017). The environment is covered by the overlapping place fields of different place cells and the population activity can be used to decode the animal’s position and even future trajectories once the firing fields have been mapped (Wilson & McNaughton, 1993; Dragoi & Tonegawa, 2011; Pfeiffer & Foster, 2013; ÓlafsdÓttir et al., 2015; Pfeiffer, 2020). In the five decades since the discovery of place cells, many other cell types supporting spatial representation have been found, including (but not limited to) head direction cells (Taube et al., 1990a,b), border cells (Solstad et al., 2008), and grid cells (Fyhn et al., 2004; Hafting et al., 2005). Especially grid cells have received much attention: Located in the medial entorhinal cortex (MEC) (Fyhn et al., 2004) and pre- and parasubiculum (Boccara et al., 2010), these cells fire at regularly spaced intervals, tiling the environment into a regular hexagonal lattice. Grid cells are dorsoventrally organized into discrete modules of constant scale and orientation; between modules, scale is increasing in discrete steps (Hafting et al., 2005; Stensola et al., 2012). Within a module, grid phase and orientation between neighboring cells remains fixed (Fyhn et al., 2007). Due to their regular firing properties and the fact that the pattern persists in complete darkness, grid cells are generally thought to be involved in path integration (Hafting et al., 2005; Fyhn et al., 2007; Rowland et al., 2016; Grieves & Jeffery, 2017).

Grid cells are a common spatially modulated cell in the MEC (Hafting et al., 2005) and form a major input to place cells (Brun et al., 2008; Jeffery, 2011; Rowland et al., 2016). However, silencing input from MEC to CA1 does not eliminate place cell activity (Brun et al., 2008; Bush et al., 2014; Brandon et al., 2014) and in rat pups, place cells form before grid cells (Langston et al., 2010; Wills et al., 2012; Bush et al., 2014). Place cells show relatively stable place fields before the hexagonal grid cell pattern is fully developed, suggesting a strong input from allothetic or external cues such as landmarks or boundaries (Bjerknes et al., 2014; Muessig et al., 2015; Moser et al., 2015; Grieves et al., 2018).

This external input causes context-dependent firing changes: When the animal moves or is moved from one compartment to another, place cell specificities may undergo complete changes to an independent pattern of firing fields. This process is known as “remapping” (Muller & Kubie, 1987; Lever et al., 2002; Leutgeb et al., 2005; Colgin et al., 2008; Julian et al., 2018): Place cells randomly rearrange their firing fields to new, uncorrelated locations, form additional firing fields or cease firing altogether. When the animal later returns or is returned to the original compartment, the place cells remap to their original pattern (Lever et al., 2002; Leutgeb et al., 2005). A given place cell therefore represents different places at different times or, more correctly, in different “spatial contexts”. Remapping also occurs in grid cells: When the context changes, the firing field locations of different modules may shift but grid scale and relative orientation remain unchanged (Fyhn et al., 2007; Cheng & Frank, 2011; Marozzi et al., 2015). Due to direct anatomical projections from MEC to the hippocampus and the finding that electrical stimulation of MEC grid cells causes place cell remapping (Kanter et al., 2017), grid cell realignment has previously been suggested as a possible candidate for triggering place cell remapping (Monaco & Abbott, 2011; Bush et al., 2014).

An important puzzle piece in understanding the remapping process is the nature of the “context change” by which it is triggered. Remapping can be observed when the animal moves to different compartments within a maze, or when the color and location of cues (Bostock et al., 1991) or the overall shape and texture of the compartment is changed (Lever et al., 2002; Leutgeb et al., 2005; Wills et al., 2005; Colgin et al., 2010; Jeffery, 2011). Non-spatial contexts may lead to a modulation of place cell firing rates (“rate remapping” (Latuske et al., 2018)), as has been shown for changes of the current task or goal (Allen et al., 2012) or in the dependence on recent history (Keinath et al., 2020). For a review of remapping types, see Latuske et al. (2018). Importantly, remapping seemingly does not occur if the context remains unchanged: For example, grid cell patterns are undisturbed within each compartment in Fyhn et al. (2007); Derdikman et al. (2009) and Carpenter et al. (2015).

This suggests that the current compartment is equivalent to the current context, for as long as no remapping occurs, the rat must be within the same area. Therefore, remapping suggests a regionalization of the rat’s cognitive map, although the rat may not have direct access to that information. In this sense, the pattern of active cells provides both information about the current context and the position of the animal within that context (Latuske et al., 2018). Overall, the findings suggest that remapping is an active process that reflects the recruitment of a unique cognitive map from a “cognitive atlas” (Julian et al., 2018), a putative higher-level hierarchical structure of spatial representation.

How can the hippocampal-entorhinal circuit acquire, maintain and reactivate a multitude of unique, context-dependent maps in a noisy infinite-context world? Some hints to this question may be hidden in a number of particularities of the remapping process:

1. Similar contexts lead to similar representations. In visually or geometrically identical but otherwise separate compartments, place and grid cell firing patterns have been observed to repeat (Derdikman et al., 2009; Spiers et al., 2015; Carpenter et al., 2015; Grieves et al., 2016; Harland et al., 2017). That is, the place and grid cell activity at corresponding positions in different rooms is highly correlated. This similarity is behaviorally relevant: In a spatial memory task by Grieves et al. (2016), rats had to memorize odor-coded buried rewards in each of four identical rooms connected by a lateral corridor. The rooms were either arranged in parallel or radially around a curved corridor (Fig. 1). The animals learned the task faster and performed significantly better in the distinguishable radial condition, where place cells remapped to different patterns in each room, but failed in the parallel condition, where the patterns repeated (Grieves et al., 2016). Remapping also does not just reflect a direction or mode of movement: Derdikman et al. (2009) only observed remapping in a hairpin maze with actual physical walls, and not in rats that were trained to run comparable zig-zag lines in an open room.
2. Entrance orientation is sufficient to distinguish contexts. The superior performance in the radial condition is attributed to the head direction system which is assumed to maintain a global reference while the animal explores the different compartments (Harland et al., 2017). If the orientation of the compartments is the same and they are entered from the same geocentric direction, the reference is also the same and the rooms are confused. On the other hand, if the compartments extend into different directions, such as in the radial arrangement in Fig. 1, the head direction reference may be used to detect this difference and the compartments can be distinguished. If the head direction system is silenced, this ability disappears and the radially arranged rooms also show the same local pattern (Harland et al., 2017). In agreement, the inverse case, where locally and globally identical compartments show different firing patterns, has also been reported (Skaggs & McNaughton, 1998; Tanila, 1999; Grieves et al., 2016; Harland et al., 2017; Keinath et al., 2020). Notably, Keinath et al. (2020) found that the most recent entrance influences the place cell firing rate in a compartment with multiple entrances, even if the firing field positions remain the same. Overall, the distinguishing cues seem to be entrance position and direction relative to the compartment, i.e., also orientation references.
3. Local cues only play a limited role. Overall, the remapping dynamics suggest a strong bias towards local cues, i.e., the sensory and geometrical information of the current surroundings. These findings led to the formulation of border or boundary vector cell models of place cell firing (Hartley et al., 2000; Barry et al., 2006; Bush et al., 2014; Grieves et al., 2018): These cells fire whenever the animal is located at a preferred distance and angle from a nearby wall and thus reflect the local geometry. Based on the repeating patterns in identical compartments, the models suggest that geometric cues measured by boundary vector cells and mediated by the head direction system are the main determinants for place fields. However, boundary vector cells show no remapping. The cells have been reported to continue firing at the same fixed distance and allocentric direction from boundaries across multiple environments, regardless of context (Lever et al., 2009). This is exemplified in e.g., Spiers et al. (2015), where the change of wall color in one of multiple identically shaped parallel rooms caused a different place cell pattern to emerge in the changed room only. Therefore, remapping dynamics must occur at least partially independent of boundary vector cells, and models based on them require additional context (e.g., “contextual gating” in Grieves et al. (2018)).
4. Place and grid cells remap together. Remapping appears to happen nearly instantaneously (Jezek et al., 2011) and both cell types remap concurrently (Fyhn et al., 2007) at the maxima of contextual difference, such as gateways between compartments or region transitions (Tanila, 1999; Leutgeb et al., 2005; Derdikman et al., 2009; Carpenter et al., 2015; Grieves et al., 2016). Therefore, the context-defining information must be available immediately at the transition. After remapping, patterns usually remain stable and no further remapping occurs as long as the local context remains the same.

**Figure 1:**
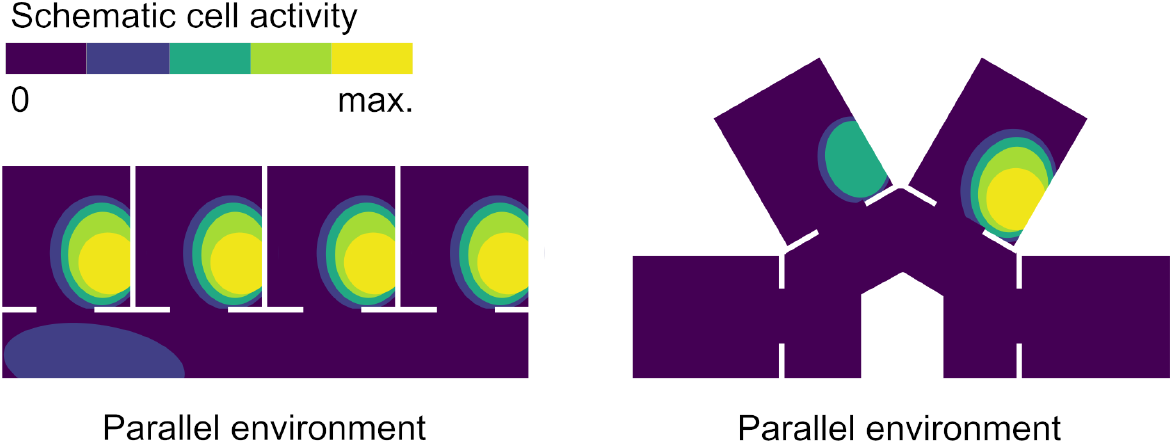
Schematic example of findings by Grieves et al. (2016). The place fields of the same place cell are shown in two environments. Firing repeats at corresponding positions in each room in the parallel environment, but not when the rooms are arranged radially.

Overall, the findings imply that gateways between compartments play a special role: Distinct place and grid cell patterns arise immediately if the compartments can be distinguished at the entrance. On the other hand, if they cannot be distinguished, the context is the same and the cell patterns reflect that similarity. In this sense, the context of the current environment would be based on the most recent gateway rather than just local cues.

Here, we propose a combined grid-and-place cell attractor model augmented with a place memory that is able to capture these remapping dynamics. The attractor is based on the observed connectivity loop of direct projections from MEC to HC (Brun et al., 2008; Jeffery, 2011; Monaco & Abbott, 2011; Rowland et al., 2016) and recurrent connections back from HC to MEC (Hafting et al., 2005; Boccara et al., 2010; Bush et al., 2014; Rennó-Costa & Tort, 2017): As in other models (Gaussier et al., 2007; de Almeida et al., 2009; Cheng & Frank, 2011; Monaco & Abbott, 2011; Lyttle et al., 2013; Li et al., 2020), the activity of multiple grid cell modules is summed to form realistic place fields; in addition, we assume that place cells also connect to grid cells and may thus support the current attractor or drive the system into a new one.

Place cell remapping means that path integration is overruled at certain points, based on local position information at that point. This implies that the recognition of local position information is essential. We specifically look at natural region transitions in the form of gateways between rooms and introduce a mechanism called the “gateway identifier”, which represents context at entrances, including head direction. Each gateway has two gateway identifiers, one on each side.

To obtain repeating place and grid cell patterns in similar environments, the model has access to a “gateway database”, a memory that stores and reactivates place cell patterns associated with the different gateway identifiers. The model exploits the ability of place cells to remap to arbitrary positions when directly stimulated (Diamantaki et al., 2018): Whenever a known gateway is passed, the associated pattern is reactivated and the place cell activity changes to it. Recurrent input from the place cells to grid cells then causes a shift in the grid cell firing fields until the matching attractor is reached.

Within a region, cell activity is based on path integration only. As such, firing fields are only determined by the local position information encountered at the last gateway and some additional noise. Importantly, entrances to ostensibly different regions may share the same gateway identifier if they share the same local position information, leading to the experimentally observed repetition of activity in similar compartments. Finally, when the system identifies an unknown entrance, it randomly remaps and the new activity pattern is added to the database.

Our model is related to models of working memory by Bouchacourt & Buschman (2019), where representations are maintained through recurring connections between a structured sensory layer (the grid cells) and a random, unstructured layer (the place cells). As a model of working memory, the model is inherently limited in its capacity once representations in the unstructured layer start to overlap. In this sense, the gateway database is a long-term memory that binds working memory patterns to context and is able to reinstantiate these patterns in a top-down manner.

In the following, we show that the model is able to replicate the remapping dynamics observed in multicompartment environments (e.g., Tanila, 1999; Fuhs et al., 2005; Carpenter et al., 2015; Grieves et al., 2016), and we propose a novel multicompartment setup (L-shaped rooms, Virtual environments) to confirm or refute the role of gateways as the main determinants for local context.

## 2. Materials and Methods

### 2.1. Model overview

The model consists of an attractor neural network with fixed weights, simulating the cognitive map of a rat exploring a series of virtual environments. The cell activity is iteratively simulated in real-time; we denote the discrete time step of a single computational sequence as *t*.

The core of the model is a grid cell attractor based on (Guanella et al., 2007). Input arrives in the form of an egomotion velocity vector, which shifts grid cell activity peaks to form the well-known hexagonal firing fields. Grid cells are arranged in multiple layers that differ in orientation and gain; from these, place cell activity is obtained by summing over the grid cells in a winnertake-all scheme. We adopted this modeling approach to place cell firing fields from previous studies (de Almeida et al., 2009; Monaco & Abbott, 2011; Lyttle et al., 2013; Li et al., 2020) in order to keep the system simple. However, local position information can easily be added as another input to the place cell layer. The place cells then feed back into the grid cell module to complete a larger attractor loop (Fig. 2).

**Figure 2:**
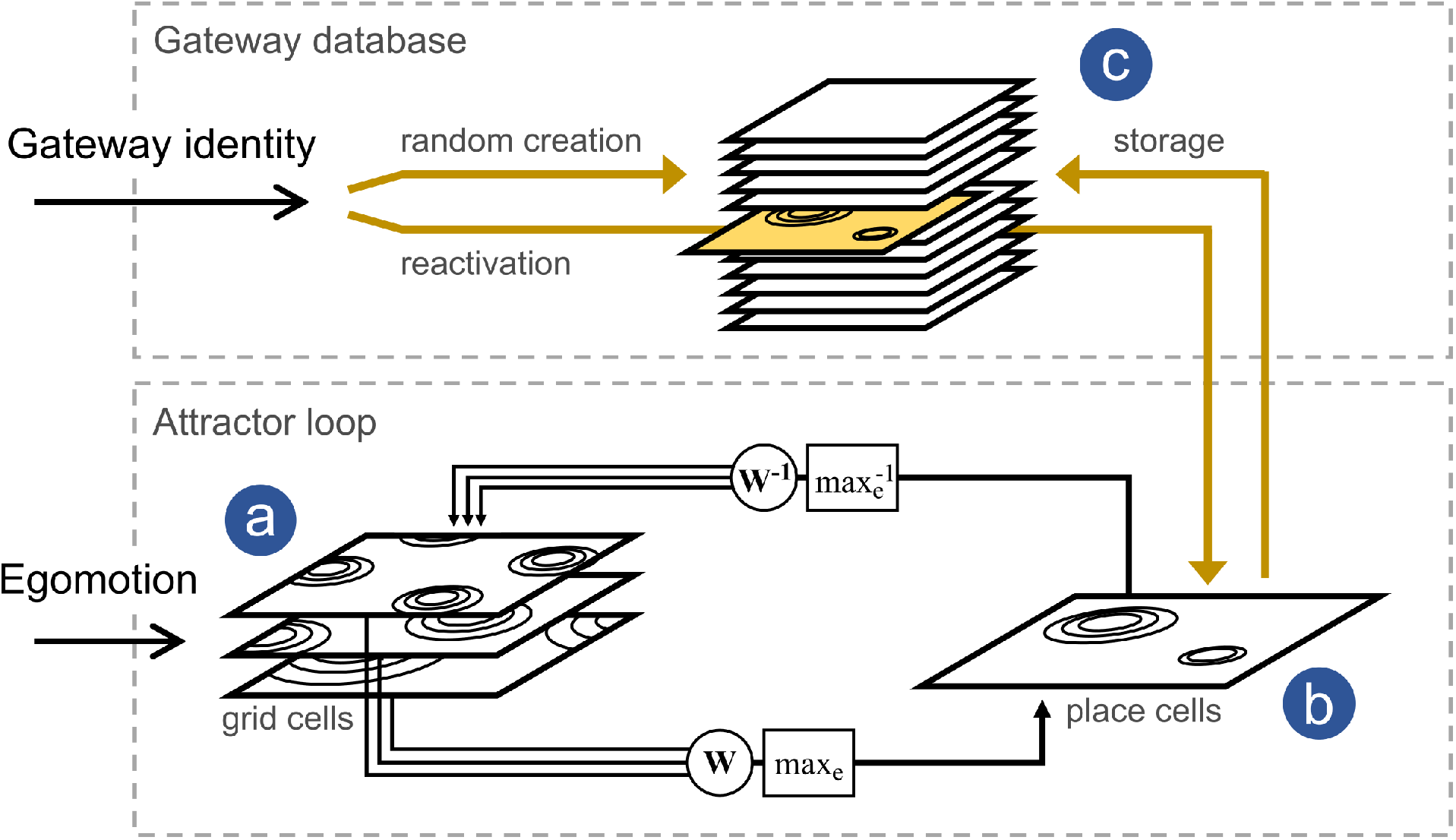
Model overview. **(a)** 3 × 100 grid cells are modeled by moving a stable activity peak over a toroidal surface. The cells form three layers with different input gains, resulting in hexagonal patterns with different scale and orientation. The circles on the layers sketch the firing fields, not the attractor peaks (which are the same for every layer). **(b)** Grid cell activity is combined in a weighted sum and filtered by an (invertible) winner-take-all scheme to form the place cell activity. The place cells in turn activate the grid cell module with the inverse operation, completing the larger attractor loop. **(c)** At region transitions such as room entrances, place cell patterns are memorized so that they may be recalled in the future (yellow arrows). When the virtual rat enters another region, place cell activity is either randomized or a corresponding place cell pattern is reactivated, followed by a shift of the grid cell firing peaks to the best matching attractor.

To explore remapping dynamics, a predefined remapping signal is triggered whenever the virtual rat passes through a gateway and the corresponding gateway identifier is activated. It causes the place cells to either remap to a new, random pattern if the gateway is unknown, or to a previously memorized pattern if the gateway is recognized. By memorizing patterns at the gateways, compartments become associated with specific cell configurations, which remain active for as long as the rat does not move to another compartment.

To distinguish rotated but otherwise identical compartments, head direction information is also implicitly included, but simply assumed to be perfect in its compass-like function. I.e., the virtual rat is always aware of “true north”. This allows us to essentially ignore the virtual rat’s heading and treat the step-wise path integration as a straight-forward *x, y*-vector addition in a fixed reference frame. Throughout the paper, we use symbolic cardinal directions: North, east, south and west mean up, right, down and left in the figures.

### 2.2. Grid Cells

Egomotion is processed by the grid cell module, consisting of 300 grid cells divided into three independent granularity layers with increasing scale and varying orientation. Three layers are required for realistic remapping based on independent shifts, as described in (Monaco & Abbott, 2011). In accordance with measurements by Stensola et al. (2012), the grid scale, i.e., the shortest distance between two activity peaks, was set to 40cm for the most fine-grained granularity layer and increased by a factor of 1.42 per layer to 57cm and 80cm for the coarser layers.

In the rodent brain, grid cells in the same granularity layer have been reported to share the same orientation (Stensola et al., 2012), are aligned to nearby walls, and the orientation remains constant across different contexts (Krupic et al., 2015). In our model, grid orientation was fixed at 8.8° for layers 1 and 2 and 98.8° for layer 3, in accordance with the measurements by Krupic et al. (2015).

Within a layer, a stable activity peak is maintained by attractor dynamics (Guanella et al., 2007). *n*^2^ = 100 cells form a topologically organized periodic sheet where neighboring cells have neighboring firing fields (Fig. 3a). The cells are connected to all others by short-range activation and long-range inhibition. In order to achieve the hexagonal geometry, the cells are arranged in a rhomboid with angles 60° and 120°, i.e., composed of two equilateral triangles. The rhomboid is the primitive cell of the hexagonal grid. The Cartesian coordinates **x**_*ij*_ of grid cell *ij* within a rhomboid with lattice constant 1*/n* are

**Figure 3:**
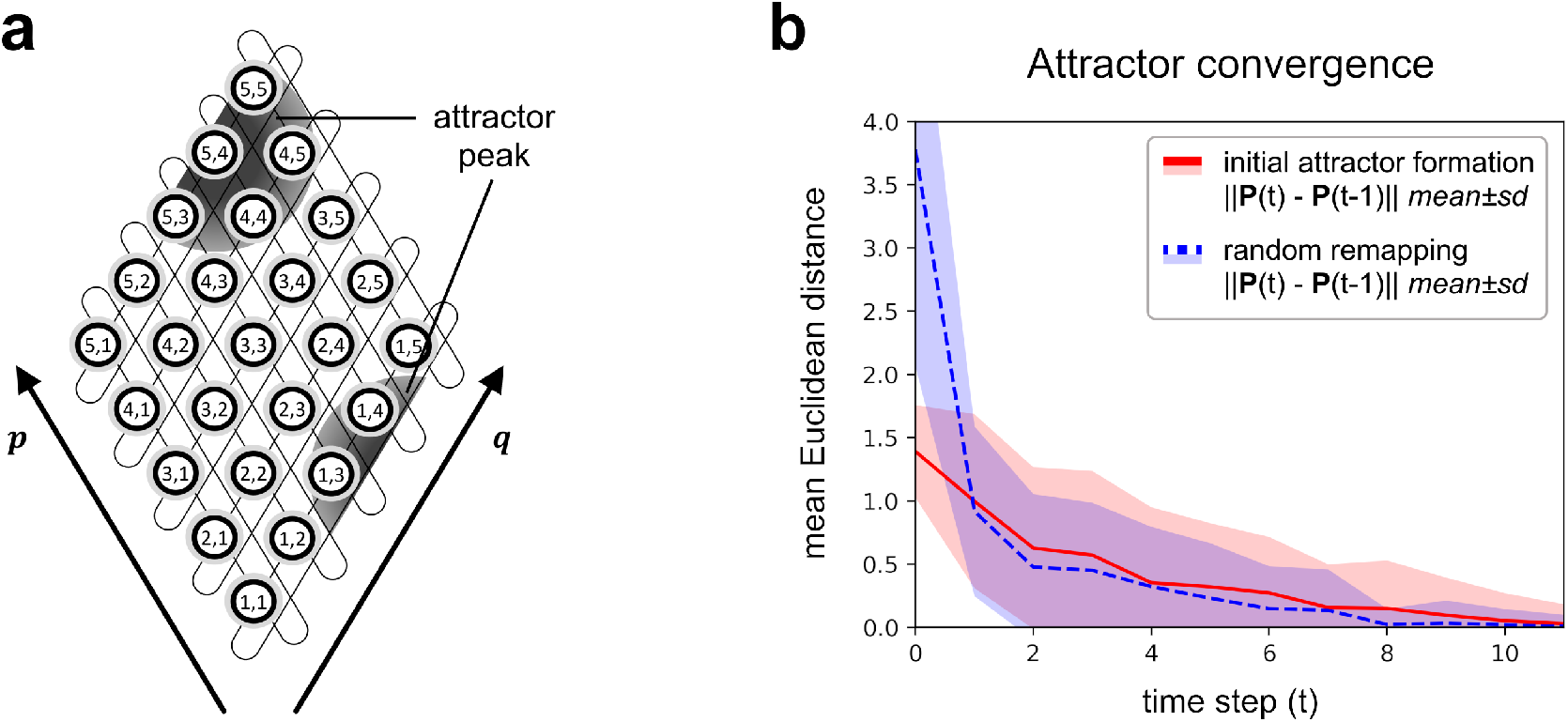
**(a)** Grid cell rhomboid (with only 5 × 5 cells for clarity). An activity peak (dark gray circle) is centered on cell *g*_5,4_. The thin lines show periodic boundary conditions along grid cell indexes *i, j*. The vectors *p, q* correspond to the side length of the rhomboid and give the shift distance *s*_*pq*_ for eq. 2. **(b)** Convergence to an attractor when the model is initialized with zero place cell activity and random grid cell activity (red), or when the model remaps to a random place cell pattern (blue). Average of 100 repetitions.

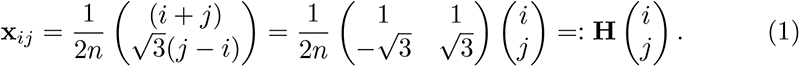

A 5 × 5 cell example is depicted in Fig. 3a. To derive an expression for the hexagonal periodic distance function, we pad a set of eight shifted rhomboids around the standard rhomboid by considering the shifts **s**_*pq*_ = **H** (*p, q*)^*⊤*^ for (*p, q*) ∈{−*n*, 0, *n*} ^2^. For given Cartesian points **x, x**′ the periodic distance function is

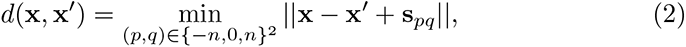

Where || *·* || is the Euclidean norm. This description of periodic boundary conditions in a rhomboid is equivalent to the twisted torus approach of Guanella et al. (2007), but can easily be generalized to an arbitrary number of grid cells.

We can now describe the activity of grid cell *ij* at time step *t*,

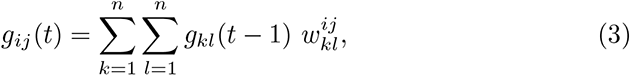

where the weights 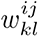 follow a Gaussian function of the periodic distance defined above,

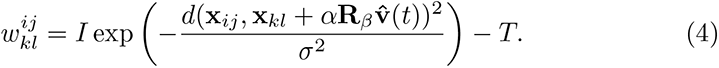

The intensity *I* = 0.3 defines the overall synaptic strength, *σ* = 0.24 is the width of the Gaussian and *T* = 0.05 is a global inhibition parameter. The scale of the grid is defined by the gain parameter *α* ∈ {0.05, 0.035, 0.025} corresponding to 40cm, 57cm and 80cm for the three layers. The orientation is given by the rotation matrix **R**_*β*_ with *β* ∈ {8.8°, 8.8°, 98.8°}, respectively. Initially, the cell activity *g*_*ij*_ (*t*_0_) is set to a random value drawn from a uniform distribution 𝒰(0, 0.1), but a stable peak is formed in the next time steps (initial attractor formation, Fig. 3b).

If the virtual rat moves with velocity **v**(*t*) = (*v*_1_, *v*_2_)^*⊤*^, synaptic weights in direction of the egomotion estimate 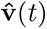 increase, causing the activity peak on the grid cell layer to move by a corresponding amount. The egomotion estimate is obtained from the true velocity by adding multiplicative noise to both components,

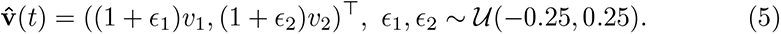

In the sequel, rather than describing the dynamics of individual cells, we combine the activities of all three layers into a single 1 × 300 grid cell activity matrix

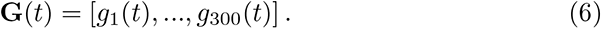

For example, the grid cell activity *g*_221_ now refers to *g*_21_ in granularity layer 3. This allows us to describe cell and layer-layer interactions in the form of matrix multiplications.

### 2.3. Place Cells

Place cells form a single layer of 300 cells. In most situations, i.e., as long as no remapping occurs, the place cell activity **P**(*t*) is obtained by summing over grid cell activity with random weights:

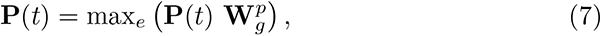

where 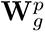 is a 300 × 300 synaptic weight matrix drawn from 𝒰 (−1, 1). The weight matrix describes the excitatory and inhibitory input from grid to place cells. As in previous studies describing place fields as originating from random summation of grid cell inputs (de Almeida et al., 2009; Monaco & Abbott, 2011; Lyttle et al., 2013), a winner-take-all nonlinearity max_*e*_ was used to sparsify place cell activity:

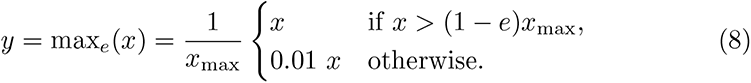

*x*_max_ is the highest *x* in the input set and *e* = 0.1 is the percentage of most active cells whose activity is maintained, while all others are attenuated by a factor 0.01.

The winner-take-all scheme restricts cell activity to local maxima, resulting in place fields qualitatively similar to real place cell recordings. Importantly, by only attenuating cells below threshold rather than setting their activity to zero, the entire operation becomes invertible, i.e., it enables place cells to predict grid cells in turn. This property is required for the combined grid-and-place cell attractor loop and remapping dynamics described in the following.

### 2.4. Combined attractor and inverse place cell - grid cell activation

By linking the place cell activity output back to the grid cells, an attractor is formed. It ensures that a specific place code will always match a specific grid cell arrangement. However, during normal exploration, the place cell input must not hinder the movement of the grid cell peak by the velocity vector. That is, the estimated activity of grid cells from place cell input, **Ĝ**(*t*) should not differ substantially from the grid cell activity **G**(*t*). The required feedback connections from the place cell layer to the grid cell layers have been shown to exist in the form of recurrent MEC-HC connections (Hafting et al., 2005; Boccara et al., 2010; Bush et al., 2014; Rennó-Costa & Tort, 2017) and could be learned by standard neural network techniques (Zhang et al., 1997; Linsker, 1997). For the purpose of this study we implemented a straight-forward inversion,

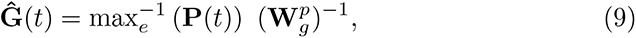

where 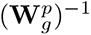 is the inverse or Moore-Penrose-pseudoinverse of 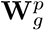 from Eq. 7. In our simulations, the random matrices came out with full rank in about 99% of the trials. The nonlinearity max 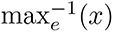 is the inverse operation of eq. 8,

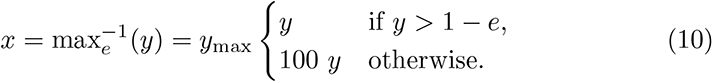

From eqs. 7 and 9, it is easy to see that **Ĝ**(*t*) ≈ **G**(*t*) except for a scaling factor. We can therefore stop the intrinsic dynamic of the grid cell module and restart it with the image of the place cell activity **G**(*t*) ≔ **Ĝ**(*t*). This will lead to a normal continuation of the grid cell dynamics, as long as **Ĝ**(*t*) falls in the basin of attraction of the current grid cell attractor. The intrinsic dynamics of the grid cell module will take care of remaining errors in the inversion process in the next few time steps. The convergence to the attractor after remapping is shown in Fig. 3b. The exact inversion of the forward weight matrix and nonlinearity is likely over-specified. The only requirement for a biological implementation would be that the inverse grid to place cell activation would fall into the same attractor basin. Since the cells are active simultaneously, it may be possible to learn the required connections via Hebbian dynamics. We opted for the matrix and nonlinearity inversion out of mathematical convenience and did not attempt to model biological neural dynamics such as synaptic plasticity or noise.

### 2.5. Remapping and gateway database

Remapping occurs whenever the virtual rat moves from one compartment to another. At the transition, the model is provided a “gateway identifier” Γ, which, in our simulations, is simply a number corresponding to the identity of the local context of the newly entered compartment. The gateway identifier is an auxiliary construct that represents the necessary information that a biological organism would need to recognize a place, e.g., from vision, olfaction or proprioception (Cheng et al., 2013). Importantly, the gateway identifier also includes head direction information, so that otherwise identical compartments may be distinguished by their orientation, but it does not include path integration or distance information, so that parallel, identical rooms cannot be distinguished by their relative position. In the virtual environments described below, entrances that look the same and are oriented in the same direction share the same gateway identifier (see Fig. 4). In the hippocampal-entorhinal network, gate information could be modeled by an additional “gate cell” connected to the place cell layer (see Discussion). This hypothetical cell would activate a specific place cell pattern whenever the room was entered through the corresponding gateway.

**Figure 4:**
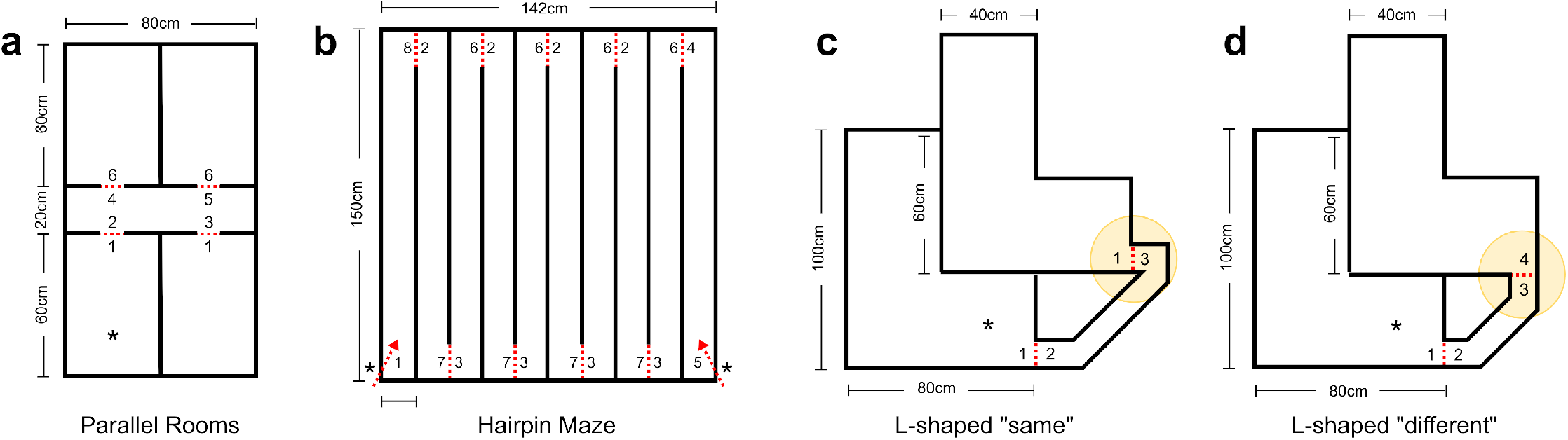
**(a) - (d)** Virtual environment floor plans used in the simulations. The red dotted lines mark region transitions and the adjacent integers the gateway identifier Γ: Equal numbers indicate the same local context and therefore cause remapping to the same patterns. The asterisk **(*)** marks the starting position. **(b)**, the hairpin maze, has two starts based on traversal direction. Note that in **(c)**, the gateway identifier of both rooms is the same, Γ = 1, but in **(d)**, it is different: Γ = 1 for the southwest and Γ = 4 for the northeast room.

In the current implementation of the model, the newly encountered gateway identifier Γ is used to index the gateway database *D* = **P**_0_, …, **P**_*n*_. If the database has an entry **P**_Γ_ ∈ *D*, the stored place cell pattern is reactivated by setting **P**(*t*) := **P**_Γ_. With the attractor rules described in eq. 9, the grid cell attractor will shift to the attractor matching the reactivated place cell pattern. As a side effect, accumulated path integrator noise is also reset when a known gateway is passed.

This mechanism may generate errors at wide gateways that can be passed with different sideways offsets. This problem could be avoided by assuming that the rat shows strict centering behavior at gateways. Here, we measure the sideways offset and add it to the displacement vector 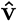 in eq. 4. In biology, offset information could for example be provided by border or boundary vector cells, or depth perception.

Resetting the place cell activity and the subsequent convergence of the network to the new grid-and-place cell attractor concludes the remapping process. Grid and place cells have completely changed their firing fields, but the change is consistent in that the same pattern will be activated whenever a given gateway is encountered. Therefore, similar compartments that share the same gateway identifiers will show the same place and grid cell patterns.

The gateway database models the rat’s overall knowledge of the compartments; it is a cognitive atlas in the sense of Julian et al. (2018) that encompasses all context-specific maps. Initially, the gateway database is empty and the model is initialized with random activity. When the rat learns a novel gateway identifier, the current place cell activity pattern is stored. This may happen in three different cases:

1. When entering an unknown room, a novel view will appear. A new gateway identifier is encountered and a random pattern of place cell activity is generated and stored in the database together with the identifier.
2. When entering a known room, the gateway identifier is recognized and the associated place cell activity pattern is loaded into the place cell layer.
3. New gateway identifiers can also be stored when leaving a room such that the room can be recognized when it is later reentered through the same gate. In order to model direction-specific firing fields observed in linear track environments (McNaughton et al., 1983; Derdikman et al., 2009), this feature of the model is applied only in wide rooms in which the rat can turn around, but not in narrow corridors.

Note that random weight matrices and random initial place cell patterns were chosen as an assumption-free approach to remapping. A more realistic approach would incorporate some sensory information about the compartment, to allow for graded similarity between contexts: Grieves et al. (2016) for example found a decrease in correlation between rotated rooms proportional to the angle between them. Another alternative to random remapping would be to continue the currently active pattern over region boundaries and show no remapping at all. In that case, grid patterns would continue undisturbed across regions.

### 2.6. Simulation and exploration procedures

For fast parallel computing of the cell matrices, the model was implemented in the machine learning interface TensorFlow (Abadi et al., 2015) running in real-time on a normal notebook PC (CPU: Intel i7-10750H, GPU: NVIDIA GeForce RTX 2070). The model was evaluated in three repetitive multicompartment environments, realized as virtual 3D models in Unity (Unity Technologies, 2021).

The 3D world was only used for collision detection and exploration, not visual input. The gateway identifiers were predefined at locations that match remapping events in experimental findings, and visually similar entrances were assigned the same identifier. All walls had the same thickness (2 cm) and height (11 cm). The environments are described in detail below.

An agent, the “virtual rat”, represented by a 4.8 cm diameter cylinder, explored the environments either by random walking (parallel rooms and L-shaped rooms) or guided by the experimenter (Hairpin maze). The virtual rat moved at a constant speed of 14 cm*/*s. For the random walk, at each time step *t*, the movement direction was rotated by a random angle drawn from u(−10°, 10°). To avoid exploration deadlocks such as continued movement into a corner, the random walk trajectory reflected off of walls. The hairpin maze environment was explored by hand to obtain complete traversals which are unlikely to result from random walking. Position data (cylinder midpoint) and cell activities were sampled once per time step, which had a duration of approximately 0.1 s.

### 2.7. Virtual environments

*Parallel rooms* (Fig. 4a). The layout is based on two- or four-room maze experiments where identical rooms are arranged in parallel and are either entered from the same direction (Skaggs & McNaughton, 1998; Fuhs et al., 2005; Spiers et al., 2015; Grieves et al., 2016) or from different directions (Tanila, 1999; Grieves et al., 2016). We combined both layouts in a single two-by-two rooms environment where two sets of parallel rooms are placed on the opposite sides of a central corridor. The rooms are locally indistinguishable except for the position and direction of the entrance. In the figure, the two rooms entered from the north have gateway identifier 1 and the rooms entered from the south gateway identifier 6. The four identifiers of the corridor are all distinguishable by the combination of view and heading.

*Hairpin maze* (Fig. 4b). We created a virtual hairpin maze similar to Derdikman et al. (2009). In hairpin mazes, cell activity has been observed to repeat every second arm and to depend on running direction (Derdikman et al., 2009). Accordingly, we defined the turns between arms as region transitions, with every second arm sharing the same gateway identifier. As described above, place cell patterns in corridors are only memorized when the corridor is entered, and not when it is left. That is, entrances and exits depend on the traversal direction, which will lead to direction-selective firing fields. The maze was traversed on a guided path from the start to the end in separate eastbound and westbound trials.

*L-shaped rooms* (Fig. 4c,d). This novel layout consisted of two adjacent identical L-shaped rooms connected by a single diagonal hallway. In the “same context” condition (Fig. 4c), both rooms are entered at the same position (south-east corner) and with the same heading. The rooms are therefore locally indistinguishable and share the same gateway identifier. In the “different contex” condition (Fig. 4d), one entrance was rotated by 90° so that the rooms are entered with a different heading. Previous studies have suggested that entry direction alone may be sufficient for rats to distinguish otherwise identical contexts (Tanila, 1999; Grieves et al., 2016; Keinath et al., 2020). However, in these experiments, entrance position and bearing remained visible from the entire room and could have been used as landmarks. Models reflecting local cues alone would therefore predict the same activity in the far part of the room in both conditions. In the L-shaped rooms, the entrance is not visible from the far part of the room. Therefore, if contextual information can be maintained in absence of context-defining cues, cell patterns or behavioral measures should differ even if the rooms are completely identical.

### 2.8. Data analysis

For each environment, place and grid cell activity maps were produced by sorting the localized activity of each cell into 2 × 2cm bins, averaged over all time steps. Activity maps were smoothed by a 3 × 3 bin Gaussian kernel. Due to the radius of the virtual rat and the wall thickness, activities in different compartments were at least three bins apart, and smoothing did not extend across walls.

To assess firing field repetition, Pearson correlations were calculated between different regions of interest on the same cell activity map. Only same-size regions were compared; we did not attempt to correlate scaled cell activities. To account for random variations, each environment was explored ten times with different random number seeds, and the correlations were averaged. Place and grid cell results are reported separately.

Additionally, as a means of confirming the regions of interest as correlation maxima, we created autocorrelograms by correlating activity maps with binwise shifted versions of themselves.

## 3. Results

In the following, we report model predictions for a series of environments, in the form of binned place and grid cell activities. In the model, the cells form a combined attractor where the activity of one cell type directly influences the other. Therefore, correlation results for the different cell types, although reported separately, are highly similar.

### 3.1. Parallel rooms

We compared place and grid cell activity of matching bins for the four rooms (numbers 1 to 4 in fig. 5a) combined into parallel (west: rooms 1 and 3 versus east: 2 and 4), opposed (north: 3 and 4 versus south: 1 and 2) and rotated opposed (north: 3 and 4, rotated 180°, versus south: 1 and 2), rotated by 180° to match entrance direction. All place cells in all repetitions showed space-dependent activity patterns. Consistent with findings from animal research (Tanila, 1999; Fuhs et al., 2005; Spiers et al., 2015; Carpenter et al., 2015; Grieves et al., 2016), we observed highly repetitive activity for the parallel condition (mean east/west correlation 0.75, 95% confidence interval (CI) [0.69, 0.79]), but not in the opposed condition (north/south 0.01, 95% CI [−0.10,0.12]) or rotated opposed condition (north rotated/south −0.01, 95% CI [−0.13,0.10]). Parallel room correlations were significantly higher than those of opposed rooms (*u* = 0.0, *p* < 0.0001, Mann-Whitney-U-Test, MWU) and rotated opposed rooms (*u* = 0.0, *p* < 0.0001, MWU).

**Figure 5:**
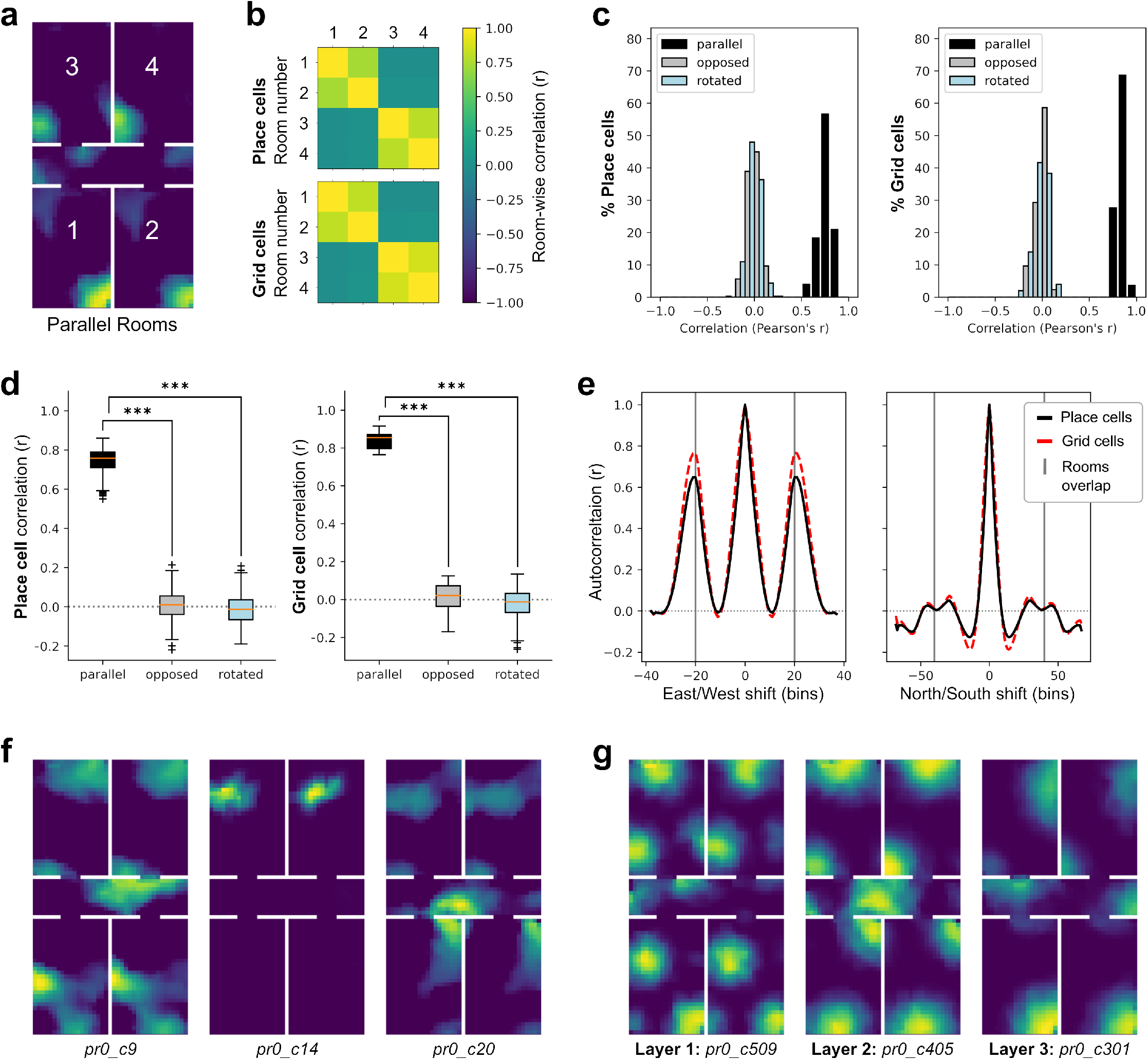
Parallel rooms. **(a)** Example of place cell activity in the different rooms (1-4) and the corridor (middle). **(b)** Average room-wise correlations for place and grid cells. As expected, the correlation between parallel rooms is high and the correlation between opposed rooms low. **(c, d)** Place cell (left) and grid cell (right) correlation distribution between parallel, opposed, and rotated opposed rooms. In agreement with the average room-wise correlation matrix in **(a)**, cell correlations cluster around 0 for the opposed condition. The distributions are relatively narrow and don’t overlap. **(e)** Linear autocorrelogram of all cells shifted in bin-wise increments, including activity in the corridor The vertical gray lines signal shifts where the rooms line up. **(f)** Examples of 3 place cells, expressing similar place fields in parallel rooms but different place fields in opposed rooms. **(g)** The same can be seen in grid cell firing fields. The grid structure in the larger layers is not visible because the rooms are shorter than the grid scale. The codes refer to the model repetition (pr0-pr9) and cell identity (0-599).

Grid cell correlations match the findings from place cells. Activity similarly repeated in the parallel condition (east/west 0.84, 95% CI [0.81, 0.87]) and was uncorrelated in the opposed condition (north/south 0.01, 95% CI [−0.10,0.12]) and rotated opposed condition (north rotated/south −0.02, 95% CI [−0.13,0.09]). Like in the place cells, the parallel room correlations were significantly higher than those of opposed rooms (*u* = 0.0, *p* < 0.0001, MWU) and rotated opposed rooms (*u* = 0.0, *p* < 0.0001, MWU), indicating that the difference in activity in the opposed rooms is not just caused by the room orientation. Overall, the results suggest that remapping at room entrances is sufficient to decorrelate cell patterns for different contexts while still producing repetitive activity under similar contexts.

### 3.2. Hairpin maze

In the hairpin maze, firing fields have been shown to depend both on the direction of movement and arm identity (Derdikman et al., 2009). Trajectories (Fig. 6a) were split into *eastbound* (from arm 1 to arm 10) and *westbound* (from arm 10 to arm 1). The agent entered the initial arm and was guided through the maze to the end of the last arm, where it exited the maze. It then reentered the same arm for a separate trajectory in the opposite direction. For each of the ten random initializations of the model, the maze was traversed twice in either direction for a total of 40 traversals.

**Figure 6:**
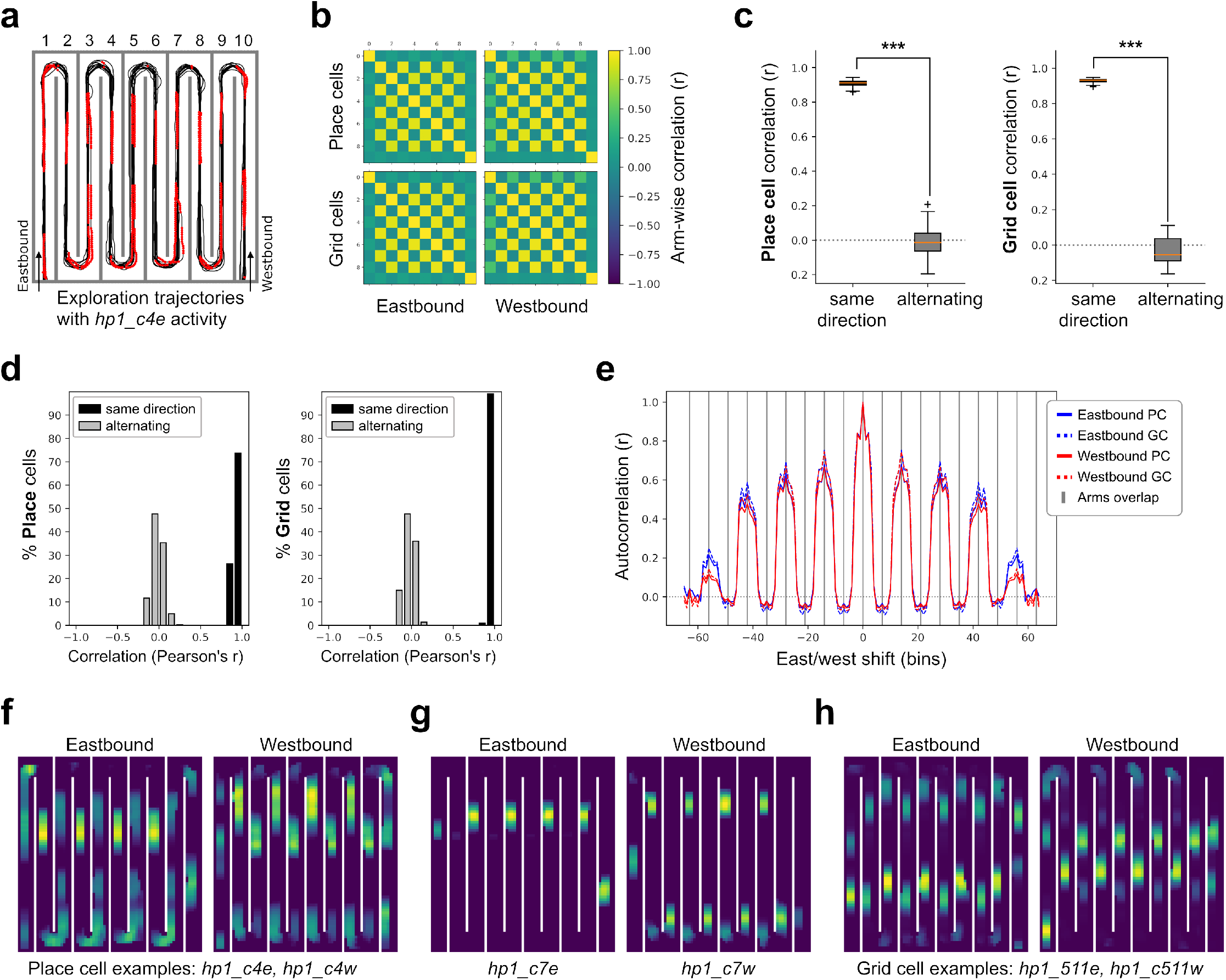
Hairpin Maze. **(a)** Exploration of the maze. The red dots mark eastbound cell activity of the cell in (f), where activity *>* 10% of max. **(b)** Arm-wise place cell (top) and grid cell (bottom) correlation for eastbound (left) and westbound (right) traversals. **(c, d)** Place cell (left) and grid cell (right) correlation distribution for the inner arms (arms 2 to 9), traversed in the same or different directions. The plots show the combined eastbound and westbound data. **(e)** Linear autocorrelogram of all cells, shifted in one-bin increments. The autocorrelation peaks whenever the activity is shifted by two arms. **(f**,**g)** Example place fields of the same cell for eastbound and westbound traversals. The cells show the characteristic repeating patterns. **(h)** Same as (f,g) but for a grid cell from the smallest layer. The grid pattern is obscured by the narrow maze arms.

Results depended on the overall traversal directions (*eastbound* vs. *west-bound*), as well as the local directions within each arm. Remapping occurred at the apices between the arms (Fig. 4b). In agreement with animal studies (Derdikman et al., 2009), model place and grid cell firing fields repeated in every second arm.

A clear checkerboard pattern (Fig. 6b) emerged from the correlation of place cells in different arms. Excluding the outer arms, arms traversed in the same direction (i.e., odd- or even-numbered arms) showed a significantly higher correlation (eastbound 0.90, 95% CI [0.88, 0.92]; westbound 0.92, 95% CI [0.90, 0.93]) than arms traversed in a different direction (eastbound −0.01, 95% CI [−0.12, 0.11]; westbound −0.02, 95% CI [−0.13, 0.09]; Same direction vs. alternating: eastbound *u* = 0.0, *p* < 0.0001, westbound *u* = 0.0, *p* < 0.0001, MWU). Firing fields in the eastbound and westbound traversals were uncorrelated (0.01, 95% CI [−0.11, 0.12]).

Grid cell results were generally similar to place cells. Grid cell activity was also highly correlated in same-direction arms (eastbound 0.92, 95% CI [0.90, 0.94]; westbound 0.94, 95% CI [0.92, 0.95]) but not in alternating arms (eastbound −0.03, 95% CI [−0.14, 0.09]; westbound −0.03, 95% CI [−0.14, 0.08]; Same direction vs. alternating: eastbound *u* = 0.0, *p* < 0.0001, westbound *u* = 0.0, *p* < 0.0001, MWU). Again, overall eastbound and westbound traversal correlations were uncorrelated (0.01, 95% CI [−0.11, 0.12]).

Correlation between the outer and inner arms was low on average (eastbound 0.14 95% CI [0.03,0.25], westbound 0.24 95% CI [0.13,0.34]). In the model, non-zero correlations between the outer and inner arms are a result of random remapping and noise, but are not indicative of any corridor similarities. The outer arms were treated as separate regions with their own gateway identifiers. In a biological animal, low but non-zero correlation might reflect a degree of similarity or confusion between the outer and inner arms, but this was not modeled.

In general, the matching correlations in the hairpin maze are higher and the confidence interval is narrower compared to the other two environments. This is likely a result of the manual exploration with frequent region transitions, which caused accumulated path integrator noise to be reset more often.

### 3.3. L-shaped rooms

The L-shaped rooms environment was explored in two entrance conditions, *same*, where both rooms were entered from the same direction, and *different*, where both rooms were entered from different directions. Note that the conditions were explored by separate, unpaired models, i.e., cells are not matched between conditions.

As in the other environments with repeating activity, place cell correlation was significantly higher in the *same* condition (0.63, 95% CI [0.56, 0.69]) compared to the *different* condition (0.06, 95% CI [−0.05, 0.17], Place cell *same* vs. *different* : *u* = 0.0, *p* < 0.0001, MWU).

For grid cells, correlations in *same* (0.74, 95% CI [0.69, 0.79]) were higher than between place cells but similarly uncorrelated in the *different* condition (0.10, 95% CI [−0.01, 0.21]) (Statistic: *u* = 0.0, *p* < 0.0001, MWU).

Firing patterns repeated when the rooms were indistinguishable at their entrances, but differed otherwise. Compared to the repeating activity conditions in the other two environments, the correlations in the *same* condition were lower. This may be a result of a less frequent path integrator reset, since the L-shaped rooms contained only two region transitions. Therefore, input noise may have played a higher role.

The bend in the rooms does not impact the results. It is not considered a gateway and causes no remapping. Because the model relies on path integration alone, it does not matter whether the entrance remains visible. The L-shaped rooms only serve as an illustration of what sort of activation to expect in a case that has not previously been covered.

## 4. Discussion

Place and grid cells have been observed to express uncorrelated firing patterns in different contexts (Muller & Kubie, 1987; Lever et al., 2002; Leutgeb et al., 2005; Fyhn et al., 2007; Colgin et al., 2008; Julian et al., 2018). The phenomenon, called “remapping”, has long been understood as a means of spatial pattern separation (Leutgeb et al., 2007) and may reflect a hierarchical organization of spatial representation (Julian et al., 2018). Remapping does not reflect purely local sensory differences, as both separate and identical patterns have been observed in similar contexts (Tanila, 1999; Carpenter et al., 2015; Grieves et al., 2016). However, the dynamics that give rise to the different firing patterns remain unknown.

In this study, we explore the idea that remapping is triggered by context change, defined as a transition between spatial regions such as the compartments of a maze. The mechanism requires a memory of contexts, implemented in our model as the gateway database. Within this memory, each context is represented as a pair consisting of (i) an identifier that can be detected from sensory input such as a snapshot and (ii) the pattern of place cell activity prevailing when the context was first encountered. Remapping occurs when a known gateway identifier is detected, e.g. by snapshot matching. In this case, the ongoing place cell activities are overridden by the stored place cell activity pattern of the recognized gateway. Projections into entorhinal cortex drive the grid cell attractor into its corresponding state and the normal path integration dynamics proceed from there.

The process is schematically illustrated in fig. 8: In a two-room environment, the agent starts by exploring the northern room. The accompanying attractor trajectory is shown as a dashed blue line in the left part of the figure. As soon as the agent enters the southern room, the gateway identifier 1 is recognized, the place cells remap to the stored pattern, and the system converges to the closest attractor, (*P*_1_, *G*_1_). Exploration of the southern room results in continuous attractor shifts until the gateway is passed again (2) and another remapping event occurs.

**Figure 7:**
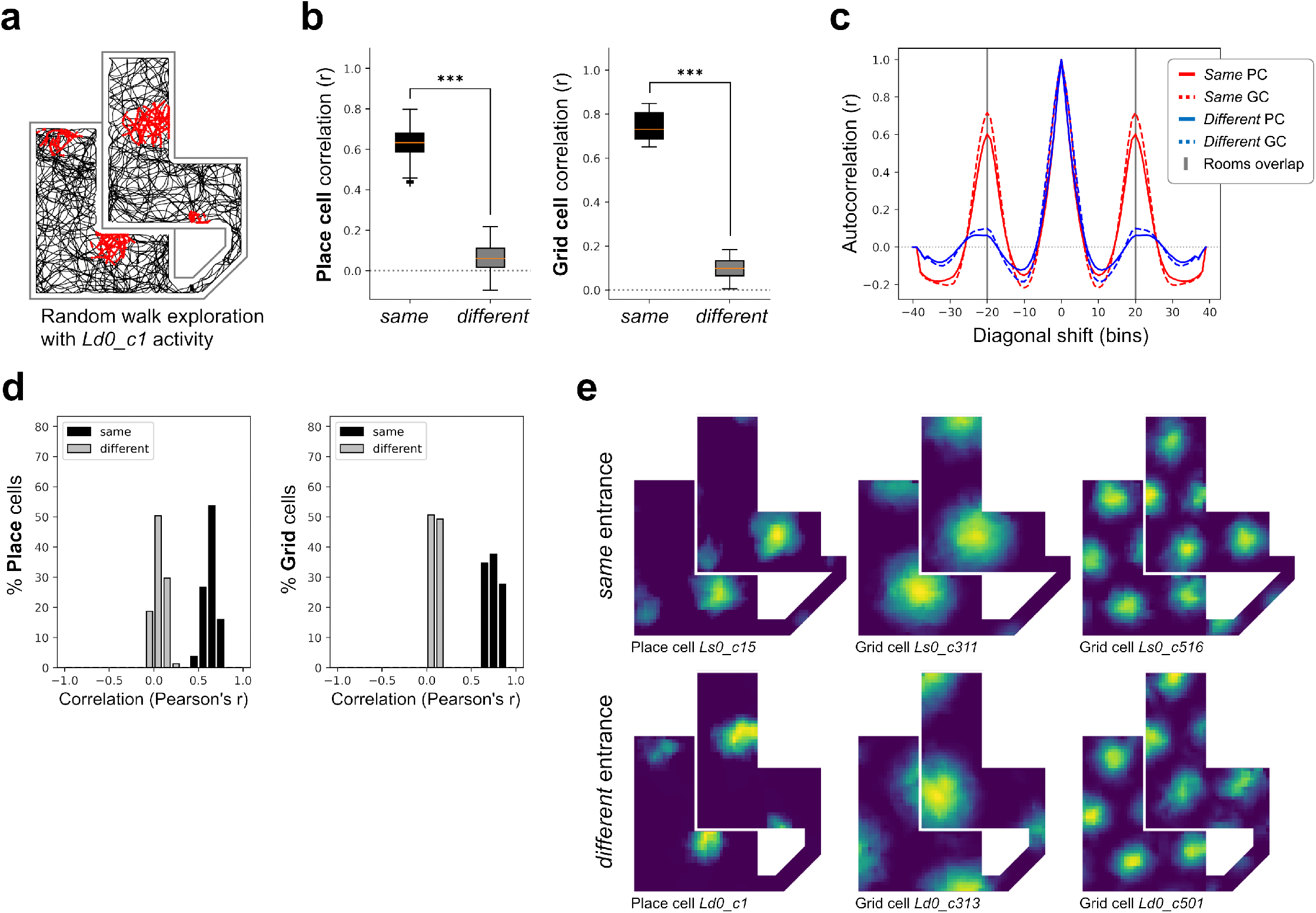
L-shaped rooms. **(a)** Random walk exploration in the *different* entrance condition. The red dots mark eastbound cell activity of the lower left cell in (e), where activity *>* 10% of max. **(b)** Place cell (left) and grid cell (right) correlation for *same* and *different* entrances. **(c)** Linear autocorrelogram of all cells, shifted diagonally in one-bin increments. The autocor-relation peaks in the *same* condition where the rooms overlap. The autocorrelogram includes firing information from the corridor, which was excluded from the other comparisons. **(d)** Cell correlations as in (b). **(e)** Firing field examples, from left to right in both conditions: Place cell, grid cell from the largest granularity layer, grid cell from the smallest granularity layer.

**Figure 8:**
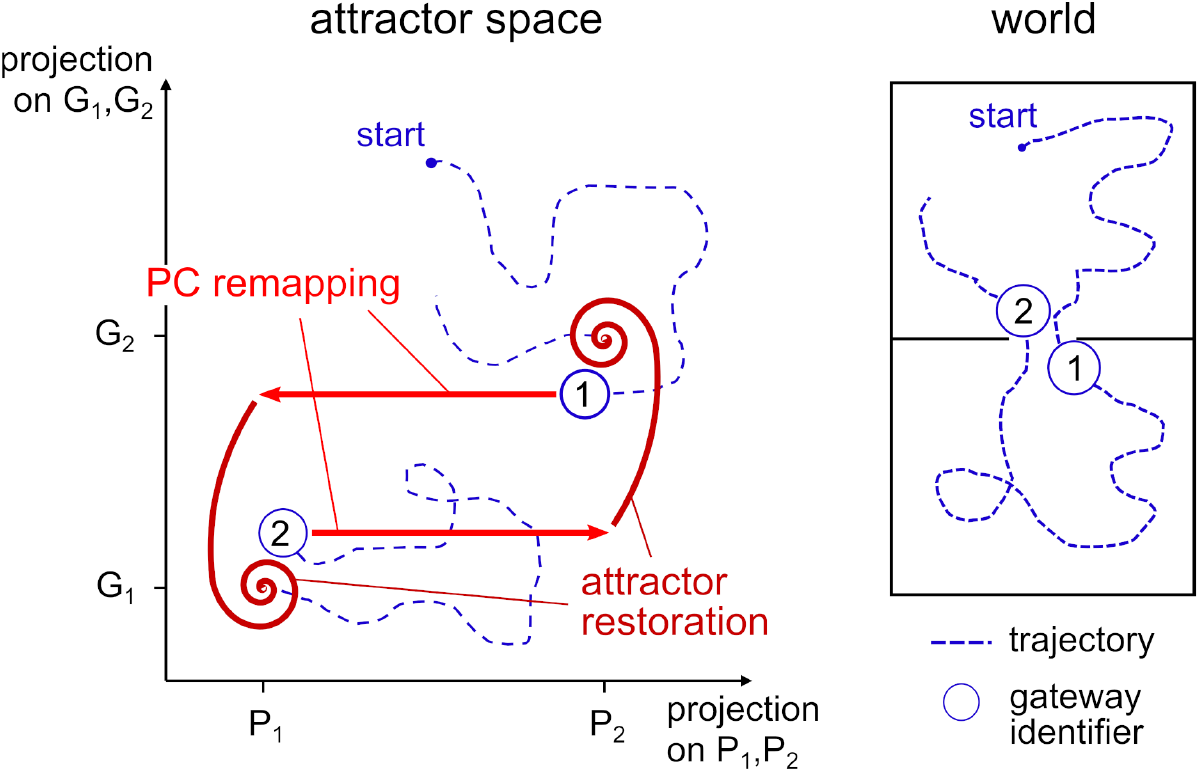
Schematic phase diagram of the remapping process. Left: 2-dimensional projection of the joint state space of place and grid cells for two attractors *P*_1_, *G*_1_ and *P*_2_, *G*_2_. The projection is spanned by the straight lines trough *P*_1_, *P*_2_ (x-axis) and trough *G*_1_, *G*_2_ (y-axis). Right: Agent trajectory in a two-room maze. For further explanation, see text.

Importantly, similar regions are assumed to share the same gateway identifier and entering them causes the same pattern to be activated. Thereby, the model was able to generate repetitive place and grid cell firing patterns qualitatively similar to those observed in multicompartment environments. In the hairpin maze, the model also replicated the directionally selective firing commonly observed in corridors.

Within each region, grid cell activity is solely controlled by noisy path integration. This causes drift over time, which is also reset by remapping. Local cues, landmarks and geometry do not influence cell activity and were indeed not modeled at all. We thus present an alternative to boundary vector cell models of place cell remapping, where geometric cues measured by boundary vector cells are the main determinants for place fields (Hartley et al., 2000; Barry et al., 2006; Bush et al., 2014; Grieves et al., 2018). In these models, repetitive patterns in similar compartments are a result of locally similar geometry.

With the L-shaped rooms environment, we propose a novel multicompartment setup where the two model types may be compared: At the far part of the rooms, the bend in the L-shape obscures the room entrance which is manipulated in two different contexts. Our model predicts that the cell patterns selected at the entrance persist over the entire region, even in parts of the rooms that are locally indistinguishable. I.e., if firing fields or behavioral measures are different between rooms, the context signal from the gateway identifier is sufficient to explain place cell remapping. If activity instead repeats, local cues are more important than the context at the entrance.

Note that we do not wish to imply that boundary vector cells play no role in place field generation. They provide position encoding independent from path integration (Bicanski & Burgess, 2020) and are able to explain place field deformation when the environment changes shape (O’Keefe & Burgess, 1996; Grieves et al., 2018) or barriers are inserted (Barry et al., 2006; Grieves et al., 2018). However, as mentioned in the introduction, boundary vector cells do not remap (Lever et al., 2009) and would need additional contextual gating to maintain distinct place cell patterns in different contexts.

### 4.1. Implications for biology

The underlying assumption behind the context-dependent cognitive map is one of hierarchy. This is directly implied by the vaguely defined term “context” itself, but also by the lack of remapping *within* a context; the spatial representation can therefore be at least divided in similar and dissimilar parts. The re-use of place cells in different population codes implies that, within a specific context, information about other contexts is unavailable. However, rats are clearly able to execute trajectories across region boundaries, so some sort of higher-level representation likely exists. Possible candidates may be cell ensembles in the medial prefrontal cortex (Hyman et al., 2012) or neurons in the perirhinal cortex (Bos et al., 2017): The former show different firing patterns depending on region, time and task and could therefore encode context (Hyman et al., 2012). The latter are neurons that fire continuously while the rat is within a specific region in space, such as an arm of a maze.

For our model, the representation of other regions is less important than the representation of transitions to these regions, i.e., the gateways. For this, we introduced the notions of “gateway identifier” and “gate cell”, which represent the recognition of the entrance and the shift to the associated cell attractor. The biological gate cell would learn a place cell pattern via neuronal plasticity and reactivate that pattern whenever the relevant stimuli occurred. Outside of the associated places, the cell should remain silent. The activation of place cells from memory is plausible: for example, with replays (Pfeiffer, 2020) and preplays (Dragoi & Tonegawa, 2011; Pfeiffer & Foster, 2013; Ó lafsdÓttir et al., 2015), the rat brain has been shown to depict sequences of place cells without the animal physically occupying the associated locations.

The gate cell attractor mechanism is intended as an alternative to contextual gating M. Hayman & Jeffery (2008), where the grid cell inputs into the hippocampus are modulated by a contextual signal to create different firing fields. Their model requires the continuous excitation and inhibition of large clusters of cells to produce orthogonal place field patterns. The advantage of our memory-driven gate cell system would be that it is more efficient: relatively few place cells need to be memorized and reactivated a single time to initially drive the attractor. I.e., the gate cell model scales much better with a high number of competing rooms or regions. On the other hand, the gate cell model cannot recover learned cell patterns at arbitrary positions, such as when the animal is placed by an experimenter (discussed below). In a sense, the distinction between the models is one of degree, i.e., how strongly the perception of context is influenced by memory and sensory information, and where exactly the neuronal connections lie.

For gateway-based context, it is of utmost importance for the animal to correctly recognize gateways, or specifically, the new region behind the gateway. A failure to do so could result in catastrophic mislocalization. Clearly, rats are able to detect changes in context, but evidence for an (over-)representation is scarce. Spiers et al. (2015) and Grieves et al. (2018) report the clustering of place fields around doorways to compartments, indicating that the doorways are at least salient landmarks; on the other hand, Duvelle et al. (2021) explicitly did not find such an overrepresentation in an experiment where the passage between compartments was blocked by lockable doors.

After detection, the rat must also be able to store gateways and associated place cell patterns in long-term memory. A possible candidate may be the retrosplenial cortex (RSC): The rat RSC is reciprocally connected to the hippocampal formation and involved in a variety of spatial tasks (Vann et al., 2009). Lesions of the RSC only cause modest spatial deficits within a context, but impair spatial memory (Vann et al., 2009) and the ability to account for context changes, such as distal cue rotation or switching from light to darkness (Wesierska et al., 2009; Pothuizen et al., 2008).

Around 9% of RSC cells are head direction cells, some of which are additionally tuned to specific movements and locations (Chen et al., 1994; Jacob et al., 2017). Here, of particular interest may be a rare type of RSC cell described in the supplementary material of Jacob et al. (2017): The cells show localized firing at the doorway between two compartments and are tuned to a particular direction. This is precisely the behavior predicted for the gateway identifier units in our model.

Mathematically speaking, the definition and detection of gateways is non-trivial. A biologically plausible solution would be the prediction or tracking of changes geometric and visual information over time; at region transitions, this change is especially high (*surprise*, Butz et al. (2004); Klukas et al. (2022)). In mobile robotics, the detection of doorways and narrow passages is a well-known problem with a variety of approaches based on cameras and laser range finders (e.g., Anguelov et al. (2004)) and posterior map segmentation (Thrun, 1998).

In humans, the cognitive map is generally thought to be hierarchically organized: in spatial memory and path planning tasks, differences arise in subjective distance estimation, recall speed (McNamara, 1986; McNamara et al., 1989) and preferred routes depending on region partition and transitions (Schick et al., 2019). In rodents, on the other hand, path planning is relatively unexplored. The best candidates are preplays, high-frequency place cell sequences observed in periods of inactivity. The preplays depict future trajectories, sometimes even through unexplored parts of the environment, indicating a planning component (Dragoi & Tonegawa, 2011; Pfeiffer & Foster, 2013; Ó lafsdÓttir et al., 2015). However, whether preplays can cross region boundaries into other sub-maps is yet unknown.

### 4.2. Remapping and place learning

Remapping at gateways is sufficient to replicate the place and grid cell dynamics observed in multicompartment environments, and it is the minimal requirement for distinct patterns in different rooms. However, there are many situations in which remapping is not quite as straight-forward. For example, what happens if the rat is enters a compartment in darkness, or if it is placed at an arbitrary position by the experimenter? What if the surrounding cues are changed while the animal is within the room? Or what about environments with multiple entrances?

Remapping can also occur if the animal does not pass through a gateway (e.g., Lever et al., 2002; Anderson & Jeffery, 2003; Colgin et al., 2010; Jezek et al., 2011; Marozzi et al., 2015). For example, in Jezek et al. (2011), rats are “teleported” between two environments by the switching of light cues, and their firing fields remap accordingly. Gateways are convenient, but in principle, arbitrary positions could be associated with a place cell pattern much in the same way. These key locations could then be recognized and reactivated whenever the animal would experience environmental contextual change, prompting it to reorient itself. In the model, this type of remapping would be straight-forward to implement, but it would come with increased memory requirements and a high risk of aliasing within a region. Remembering every single location seems plausible for small, well known environments as in Jezek et al. (2011), especially if the animals are rewarded with randomly strewn food, but it seems unlikely that a rat could immediately localize itself when placed in an open field. Alternatively, the distance and direction to a remembered place could be inferred from depth perception or path integration (i.e., information provided by the boundary vector cells (Lever et al., 2009)) and the animal may be able adjust the shift the cell patterns to the correct position. In both cases, it would be more efficient to primarily memorize salient and behaviorally relevant locations, such as gateways.

Similarly, compartments with multiple entrances are no problem if the place cell pattern is memorized when leaving through each of them, meaning that the environment has been explored adequately. Keinath et al. (2020) report a difference in firing rate but not place field position depending on the most recent entrance, i.e., the animals were able to recover the overall place code in a familiar environment. Still, there is a qualitative difference between remapping to a novel pattern on first entry and reproducing a previous pattern. The former requires some sort of gateway representation, and the latter a memory. How representations evolve in an unfamiliar environment with multiple rooms and entrances remains to be explored.

In the behavioral experiment by Grieves et al. (2016), rats eventually improved in their ability to distinguish multiple identical rooms over days, although they never reached the performance of the control group. Similarly, Carpenter et al. (2015) showed that an initially repeating grid cell pattern between two identical, parallel rooms changed into one continuous pattern over both rooms after prolonged experience. The findings indicate that the identical room problem is not unsurmountable, even if it takes a lot of time. Interestingly, one interpretation of the findings by Carpenter et al. (2015) is that, instead of learning to differentiate the two rooms as different regions, the animals combined them into one larger region with no further remapping at the gateways. This explanation nicely fits to the notion of gateways based on surprise (e.g., Butz et al., 2004; Klukas et al., 2022). After all, the entrances are not surprising if the animals know the rooms in and out. Our model currently does not simulate long-term learning dynamics and changing gateway and region definitions - it doesn’t even detect and recognize gateways automatically. However, the addition of such functions in future iterations is feasible.

### 4.3. Remapping without the MEC

Evidence points towards grid cells in the MEC as an important driving force behind hippocampal place cell remapping (Monaco & Abbott, 2011; Bush et al., 2014; Kanter et al., 2017): Grid cell remap concurrently with place cells, the MEC directly projects to the hippocampus (Fyhn et al., 2007), and electrical stimulation of the area leads to hippocampal remapping (Kanter et al., 2017). In accordance, place cell activity and remapping in our model directly arise from grid cell input. However, place cells must be at least partially independent from MEC input: Their activity persists and the cells continue to remap in different contexts, even if grid cell firing is disrupted (Brandon et al., 2014) or the entire MEC is lesioned (Schlesiger et al., 2018).

In our model, place cell activity is sustained by grid cell input and would therefore immediately cease if the input was severed. To maintain activity in the absence of grid cell input, the model place cells would either need to be augmented with recurrent weights or input from another source. This presents an opportunity to reconcile our model with boundary vector cell models (Hartley et al., 2000; Barry et al., 2006; Bush et al., 2014; Grieves et al., 2018), where local geometry, i.e., the distance to walls, determines place fields. A combined model could continue to function with interrupted path integration. Perhaps, a combined model with context-dependent grid cells and context-independent boundary vector cells could also explain the phenomenon of *partial* remapping Latuske et al. (2018), where only some firing fields remap, while others stay at the same location.

Interestingly, place fields that are newly formed while grid cells are interrupted, remain the same when grid cell activity is later restored (Brandon et al., 2014). This is consistent with the proposed feedback connections from place cells to grid cells in our model; once grid cell activity is restored, the newly formed place fields should move the grid cell activity to the closest attractor.

In conclusion, we suggest that remapping is a mechanism subserving the integration of the systems for path integration and place recognition, i.e., O’Keefe and Nadel’s (1978) Locale and Taxon systems. The cognitive map is thereby structured into a set of contexts or local charts, memorized by gateway identifiers. Within the charts, navigation is based on local metrics. On a coarser scale, the map is likely organized by the adjacencies of the charts, i.e., some sort of graph structure.

## Acknowledgment

This work was carried out at the Department of Biology of the University of Tübingen, Tübingen, Germany. This research did not receive any specific grant from funding agencies in the public, commercial, or non-profit sectors.

